# AMP-activated protein kinase (AMPK) is essential for tendon homeostasis and prevents premature senescence and ectopic calcification

**DOI:** 10.1101/2025.01.31.635920

**Authors:** LeeAnn A. Hold, Nicole Migotsky, Syeda N. Lamia, Steph S. Steltzer, Sydney Grossman, Jessica Chen, Seung-Ho Bae, Paige Cordts, Tessa Phillips, Matthew J. O’Meara, Carol Davis, Susan V. Brooks, Moeed Akbar, Neal L. Millar, Megan L. Killian, Adam C. Abraham

## Abstract

Tendinopathy is a debilitating tendon disorder affecting millions of people, characterized by pain, swelling, and diminished biomechanical properties. While the precise mechanisms underlying tendon homeostasis remain unclear, metabolic regulation plays a critical role. In this study, we combine transcriptomic analysis of human tendinopathic samples with a conditional mouse model in which Prkaa1 (encoding AMPKα1) is selectively deleted in tendon progenitors to elucidate the role of AMPK signaling in tendon homeostasis. RNA sequencing of diseased human tendons revealed downregulation of key metabolic genes, including several involved in the mitochondrial electron transport chain and AMPK signaling pathways, alongside an increase in markers associated with senescence and a secretory inflammatory profile. In parallel, mice with loss of Prkaa1 function exhibited normal postnatal development; however, by one month of age, tendons demonstrated widespread transcriptional alterations related to cell cycle regulation and ECM organization. By three months, AMPKα1-deficient tendons showed significant reductions in mechanical strength and increased expression of senescence markers p21 and p16, progressing to prominent ectopic calcification with age. In vitro studies further confirmed that tendon fibroblasts lacking AMPKα1 have altered ECM substrate adhesion profiles. Importantly, voluntary exercise partially rescued these deficits by enhancing ECM organization and reducing senescence marker expression. Collectively, our findings demonstrate that AMPKα1 is critical for maintaining energy balance, regulating ECM remodeling, and preventing premature cellular senescence in tendon. These insights highlight AMPK signaling as a promising therapeutic target and underscore the beneficial role of exercise in mitigating tendinopathic changes.

## INTRODUCTION

Tendons are poorly vascularized tissues rich in extracellular matrix (ECM) that transmit force from muscle to bone. The mechanical demand placed on these tissues requires a precise balance of metabolic processes to maintain functional homeostasis^1,2^. Tendinopathy is a tendon disorder and while the precise causes and characteristics remain to be elucidated, it is associated with results in pain, swelling, and diminished biomechanical properties. is a clinical problem that affects ∼3.5 million people in the US^3^. Risk factors for tendinopathy include age, metabolic disorders and limited exercise^3^. Tendinopathy is caused by failure of tendon to self-repair and is characterized by degenerative ECM, decreased cell viability, and poor biomechanical function^4^. Currently, treatment for tendinopathy has limited non-surgical treatment options^3^, therefore elucidating targets for a druggable therapy will improve current clinical limitations. Previously, Wu et. al established a promising molecular mechanism for treatment of diabetic induced tendinopathy through the AMP-activated protein kinase (AMPK) pathway^5^. The AMPK pathway is an intracellular stress response activated in times of nutrient deficiency, and exercise to regulate energy balance and maintain homeostasis^6^. In response to a decrease in the ATP:AMP ratio, AMPK is phosphorylated and redirects metabolism to stimulate catabolic processes while simultaneously decreasing anabolic processes^6^. While AMPK is a critical regulator of homeostasis in other tissues^7,8^ as well as a potential regulator of ECM remodeling in musculoskeletal disease^9^, little is known about how AMPK regulates energy balance and homeostasis in tendon. A key pathological characteristic of tendinopathy includes dysregulation of homeostasis, such as matrix disorganization and decreased cell viability^3^. Additionally, regulation of metabolic mechanisms play a role in tendon healing and degeneration^10,11^ establishing the potential for metabolic modulators as therapeutic targets for tendinopathy. Therefore, the potential in targeting AMPK motivates the need to define the role of AMPK in tendon homeostasis.

The basal metabolic rate of tendon is relatively low compared to other musculoskeletal tissues, possibly related to the high number of quiescent cells present in tendon during homeostasis^12^. The quiescent phenotype presents during times of limited nutrients and is coupled with a reversable cell cycle arrest. It is believed that tendon fibroblasts (TFs) become predominantly quiescent in the early stages of maturation^13^. Grinstein et. al showed in mice that after post-natal day 30 mitotic activity declines significantly and few cells continue to divide through the remainder of life^13^. Regulation of quiescence through AMPK has been shown to be age dependent in muscle stem cells^14,15^. With age, decreases in AMPK activity results in a push towards senescence through nuclear translocation of cyclin inhibitors^14^. Furthermore, activation of AMPK restored cell survival, through upregulation of autophagy and reduced nuclear translocation of cyclin inhibitors, in aged skeletal muscle stem cells^14^. These trends may be consistent in tendon, and in a study by Dai et. al, it was shown that that activation of the AMPK signaling cascade through metformin reduced the effects of aging *in vivo* and *in vitro*^16^. Using tendon progenitor cells *in vitro*, metformin reduced senescence markers p16 and p21^16^. Additionally, they showed metformin reduced the aging phenotype by inhibiting senescence marker p16 and the formation of heterotopic ossification in 10 month-old rats ^16^. Taken together, these data implicate AMPK as a possible target to regulate cellular fate in tendon and prevent age-associated functional decline.

AMPK is a heterotrimeric complex composed of α, β and γ subunits^6^. There are two α subunits, α1 and α2: AMPKα1 is ubiquitously expressed in all tissues while AMPKα2 is dominant in skeletal muscle, heart, and liver^17,18^. Reduction of AMPKα1 has been associated with musculoskeletal diseases such as osteoarthritis (OA)^19^. For example, loss of AMPKα1 leads to accelerated progression and increased severity of OA^9,19^. However, while dysregulation of AMPKα1 in some degenerative musculoskeletal diseases is known, the mechanisms that underlie its role in tendinopathy remain unclear.

In this study, we identified that the AMPK signaling pathway is transcriptionally suppressed in patients with tendinopathy. Additionally, we found that AMPK is necessary for tendon homeostasis by mediating expression of *Prkaa1* in TFs. We used Cre-lox to target deletion of *Prkaa1*, the gene that encodes AMPKα1, in tendon progenitors in mice. This targeted deletion led to normal development of tendons; however, we saw dysregulation of cell cycle and ECM biological processes in the transcriptome at 1 month of age (1M). By 3 months of age (3M), we found that tendons lacking *Prkaa1* had impaired mechanical properties and upregulation of senescence associated marker, p21. With increased age, tendons lacking *Prkaa1* developed expansive ectopic calcification and had elevated markers of senescence associated p16 and p21. Voluntary wheel running was beneficial in tendons of *Prkaa1*-deficient mice and led to increased ECM organization and suppressed p16 and p21. Our results demonstrate AMPKα1 is necessary for tendon homeostasis and exercise, an energy demanding process, which can delay senescence onset independent of AMPK α1 suggesting energy balance is crucial for cellular viability in tendon.

## EXPERIMENTAL MODEL AND STUDY PARTICIPANT DETAILS

### Human

Late-stage RC (supraspinatus) tendon samples and control hamstring (semitendinosus) tendon samples were provided by the School of Infection and Immunity at the University of Glasgow. Informed consent was obtained from participants, and the use of tissue for research was approved through the West of Scotland Research Ethics Service (REC 16/WS/0207).

### Animal Models

All procedures were approved by the Institutional Animal Care and Use Committees at the University of Michigan. ScxCre; Prkaa1^fl/fl^ male mice were bred with Prkaa1^fl/fl^ female mice to generate Prkaa1^Scx-Cre^ (cKO). Prkaa1^fl/fl^ (WT) and Prkaa1^Scx-Cre^ cKO (cKO) mice were generated for evaluation of Achilles tendon structure and function at 1-, 3-, and 9-months (1M, 3M, 9M). All mice were kept in standard housing conditions with ad libitum access to water and chow throughout the experiment.

## METHOD DETAILS

### RNA isolation and sequencing of human tendon

Tissues were collected and stored in RNALater (ThermoFisher Scientific, Waltham, MA, USA), these were subsequently physically disrupted using a ball mill homogenizer. RNA extraction was performed using RNeasy Mini Fibrous Tissue Kit (Qiagen, Hilden, Germany) with on-column deoxyribonuclease I treatment according to manufacturer’s instructions. RNA quality and quantity was obtained using Qubit fluorometer and the RNA Pico kit and Bioanalyzer 2100 system.

The NEBNext Ultra II RNA Library Preparation for Illumina Kit (New England Biolabs, Ipswich, MA, USA) was used to prepare the samples for sequencing following the manufacturer protocol. Dual Index Primers were used in the NEXNext Ultra II RNA Library Preparation for Illumina protocols. Library qualities were confirmed using Qubit fluorometer and high sensitivity DNA kit and Bioanalyzer. Samples were then sequenced using an Illumina NextSeq500 Instrument (at University of Glasgow Polyomics Facility (Glasgow Polyomics, University of Glasgow).

Differential gene expression was determined from count matrices with a paired design (e.g. Healthy vs. Diseased) in DESeq2 in R/Bioconductor (The R Foundation, Vienna, Austria) ^20^.We used the Database for Annotation, Visualization, and Integrated Discovery (DAVID) (National Institutes of Health, Bethesda, MD, USA) to analyze biological processes and Kyoto Encyclopedia of Genes and Genomes (KEGG) pathways^21,22^.

### Wheel Running

At 1M of age WT and Prkaa1^Scx-Cre^ cKO mice were randomly assigned to either remain at cage activity (CA) or were singly housed with voluntary access to running wheels (WR) for 8 weeks in standard mouse cages. Daily wheel rotations were collected and recorded. One week prior to euthanasia mice were fasted for 6hr to collect fasting blood glucose (FBG) measurements.

### Biomechanical testing of Achilles tendon

At time of euthanasia, one hindlimb from each mouse was immediately frozen at -20°C. Simple randomization of samples prior to Achille’s tendon dissection was performed and tester was blinded to all experimental groups. 24 hours before the start of mechanical testing, limbs were thawed at 4°C and prepared for photogrammetry and biomechanical testing. Both the plantaris tendon and muscle were carefully removed and the Achilles tendons and calcaneus were left intact. Cross-sectional areas (CSA) of the Achilles tendon were measured using photogrammetry with a custom pinch clamp holder attached to a motor controller (Arduino, Ivrea, Italy). With the muscle and Achilles hanging free of the pinch clamp, at least 50 consecutive images of the pinch clamped tendon were acquired using a 12mm focal length lens (Basler Fujinon Lens, Ahrensburg, Germany). Using Metashape software (Agisoft, St. Petersburg, Russia), images were aligned and converted first to a sparse point then to a dense point cloud, to create an STL surface mesh of the tendon. CSA was defined as the smallest area from the STL surface mesh generated from the Achilles tendon and was measured using a slice analysis tool in Dragonfly (Comet Technologies Canada Inc., Montreal, Canada).

A custom 3D printed fixture secured the calcaneus into place (FormLabs 3B, Somerville, MA, USA), and the proximal Achilles tendon was clamped in a textured grip and screwed into place (Imada, Northbrook, IL, USA). The assembled grip with the secured Achilles tendon was placed into a phosphate buffered solution (PBS) bath at 37°C using a temperature controller (MA160, Biomomentum Inc., Laval, Quebec, Canada) and secured with a pin to a tensile testing frame with a multi-axis load cell (±70 N; Mach-1 VS500CST, Biomomentum, Laval, Quebec, Canada). Samples were preloaded to 0.1 N, and gauge length was measured as the distance between the secured calcaneus and the textured grip.

The Achilles tendons were then preconditioned for 10 cycles (±0.1 N at 0.1mm/sec) followed by the load to failure test at 0.1mm/sec. Off-axis load (forces in X and Y, torques in X, Y, Z) were collected to assess off-axis loading for the duration of the experiment. Instantaneous grip-to-grip strain was calculated as the displacement divided by the original gauge length, *L*_0_. For the 1M, 3M and 9M WT and cKO age-dependent tests maximum load, maximum stress, and maximum strain were calculated using a custom MATLAB script (R2017 or later, MathWorks, Natick, USA). Data between cKO and WT Achilles tendons at 1M, 3M and 9M were compared using a two-way ANOVA with post-hoc Sidak’s multiple comparisons to identify specific differences (Genotype, Time).

### RNA isolation and sequencing of murine tendon

1M and 3M old as well as WR and CA WT and Prkaa1^Scx-Cre^ cKO mice were euthanized and Achilles tendons were immediately dissected under ribonuclease (RNase)–free conditions and flash frozen (*n* = 3/group). Tissues were mechanically pulverized in TRIzol, and total RNA was isolated using spin-columns (with on column genomic DNA digestion. RNA Integrity Number (RIN) was measured using Bioanalyzer (Agilent, Santa Clara, CA, USA). This pool was subjected to 151bp paired-end sequencing according to the manufacturer’s protocol (Illumina NovaSeqXPlus). BCL Convert Conversion Software v4.0 (Illumina, San Diego, CA, USA) was used to generate de-multiplexed Fastq files. Reads were trimmed using Cutadapt v4.8^23^ and FastQC v0.11.8 was used to ensure the quality of data^24^. Next, Fastq Screen v0.15.3 was used to screen for various types of contamination. Reads were then mapped to the reference genome GRCm38 (ENSEMBL), using STAR v2.7.8a ^25^ and assigned count estimates to genes with RSEM v1.3.3 ^26^. Alignment options followed ENCODE standards for RNA-seq. Multiqc v1.20 compiled the results from several of these tools and provided a detailed and comprehensive quality control report ^27^. Differential gene expression was determined from count matrices with a paired design (e.g., WT v cKO, WT WR v. CA or cKO WR v. CA) in DESeq2 in R/Bioconductor^20^. We used the Database for Annotation, Visualization, and Integrated Discovery (DAVID) to analyze biological processes and Kyoto Encyclopedia of Genes and Genomes (KEGG) pathways^21,22^.

### Histology Preparation and Imaging

Distal hindlimbs from WT and cKO tendons (n = 3 per sex/group) were dissected, fixed in 4% paraformaldehyde (PFA), and decalcified in 14% ethylenediaminetetraacetic acid (EDTA) and processed for paraffin embedding. Tissues were sectioned at 5 μm in the sagittal plane and stained with either Hematoxylin and Eosin (H&E) (StatLabs, McKinney, TX, USA), Silver Nitrate, Picrosirius Red (PSR) (StatLabs, McKinney, TX, USA) or processed for immunohistochemistry (IHC), and cover-slipped with an acrylic mounting media. H&E, Silver Nitrate (FisherScientific, Waltham, MA, USA) and (IHC) slides were imaged on a bright-field microscope (ECLIPSE Ni-U, Nikon) at 10x and analyzed using QuPath. PSR slides were imaged with a 10x objective on an epi-fluorescent microscope (dmi600b, Leica, Wetzlar, Germany) and analyzed using quantitative polarized light imaging (qPLI) analysis.

### Immunohistochemistry

For immunohistochemistry, paraffin sections were deparaffinized followed by heat-mediated antigen retrieval with an Antigen Unmasking Solution (VectorLabs, Newark, CA, USA), Citrate-Based or Tris-EDTA (pH 9.0) for 2hrs at 65°C and blocking (2.5% horse serum). Slides were then incubated in primary P16-INK4A (ProteinTech, Rosemont, IL, USA) and p21 (ProteinTech, Rosemont, IL, USA) antibodies overnight at 4°C, followed by washing and incubation with appropriate secondary antibodies. ImmPACT DAB Substrate Kit, Peroxidase (HRP) (VectorLabs, Newark, CA, USA) was used for detection, and slides were counterstained with hematoxylin and cover-slipped with Shandon Mount.

### qPLI Analysis

The degree of linear polarization (DoLP) and angle of linear polarization (AOP) images were acquired using a polarization camera and a circular polarizing lens. The mean DoLP and standard deviation of the AOP were analyzed using Math and SciPy Stats libraries in MATLAB (Natick, MA, USA). Data from cKO mice were compared to WT controls at 1M, 3M and 9M using a two-way ANOVA with post-hoc Sidak’s multiple comparisons with Graphpad Prism (San Diego, CA, USA) to identify specific differences (Genotype, Time).

### FIJI Analysis

Silver Nitrate slides were imaged and collected for percent lesion area analysis by FIJI (NIH, Bethesda, MD). First, the Achilles tendon region of interest area was selected using morphological features to determine total area. Next, using the polygon shape tool, the positive lesion was outlined, and the area was measured. All measurements were taken three times to ensure reproducibility. Percent lesion area was calculated by dividing lesion area(s) by the whole tendon area. Data from Prkaa1^Scx-Cre^ cKO mice were compared to WT controls at 9M using an unpaired two-tailed t-test.

### QuPath Analysis

IHC slides were imaged and collected for percent positive analysis by QuPath^28^, an open-source software for digital pathology. First, the regions of interest in Achilles tendons were selected using morphological features and areas were measured. Next, color deconvolution was performed by QuPath to digitally separate stains (Hematoxylin and DAB) within each tissue sample. Positive cell detection, highlighting DAB positive cells, was then performed to differentiate cells. Total cell detection was collected from counting hematoxylin positive nuclei. Data from Prkaa1^Scx-Cre^ cKO mice were compared to WT controls at 3M and 9M using an Ordinary two-way ANOVA with uncorrected Fisher’s LSD, with a single pooled variance to identify specific differences (Genotype, Time).

### Nuclear Circularity

Slides were deparaffinized and stained with antifade mountant with NucBlue. Slides were imaged with widefield fluorescence on a Cytation 10 (Agilent, Santa Clara, CA, USA) at 20x. Nuclei counts and circularity were calculated with the Gen5 (Agilent, Santa Clara, CA, USA) cellular analysis tool. Data from Prkaa1^Scx-Cre^ cKO mice were compared to WT controls using a one-way ANOVA with Tukey’s multiple comparisons test with a single pool variance.

### Cell Isolation and Culture

Prkaa1^Scx-Cre^ cKO mice were euthanized and tails were removed. Skin was removed from tail and tendon fascicles were carefully dissected and placed in a 60mm petri dish with Dulbecco’s Modified Eagle’s Medium/Nutrient Mixture F-12 (DMEM: F12), 10 % Fetal Bobine Serum (FBS) and 1% Penicillin/Streptomycin (P/S), media was changed every other day. Tail TFs were allowed to “crawl out” for two weeks before passaging for β-Galactosidase staining and ECM arrays.

### β-Galactosidase (Beta-Galactosidase) Staining

Prkaa1^Scx-Cre^ cKO Tail TFs were passaged into a 12 well plate in DMEM: F12, 10% FBS and 1% P/S for 24h. Media was changed to DMEM: F12, 1% FBS and 1% P/S for 96h, media was changed every other day. Cells were fixed and stained per manufacturer’s instructions with the Senescence β-Galactosidase Staining Kit (Cell Signaling, Danvers, MA, USA) and imaged on a Lionheart RX digital microscope (Agilent, Santa Clara, CA, USA).

### ECM Arrays

Prkaa1^Scx-Cre^ cKO Tail TFs were passaged onto a per manufacturer’s instructions in DMEM: F12, 10% FBS and 1% P/S for 24h. After 24h, cells were fixed with 4% PFA and, rinsed with PBS stained with Hoechst. Slides were imaged using a Leica THUNDER (, Leica, Wetzlar, Germany) and nuclei were counted using FIJI (National Institutes of Health, Bethesda, MD, USA) with the Stardist plugin^29^. Counts were collected and analyzed with Bayesian statistical modeling.

### Bayesian Statistical Modeling

Nuclear Counts for the WT and Prkaa1^Scx-Cre^ cKO Tail TFs on different substrates were compared with the fit of a range of two Bayesian regression models, baseline and interaction. In the baseline model the cell counts are assumed to occur assumed to occur independently at random only. With the interaction model, the cell counts are dependent on the genotype and substrate.

## QUANTIFICATION AND STATISTICAL ANALYSIS

For graphs in figures 2,4 and 5 data are shown as mean ± standard deviation (SD). GraphPad (GP) p-value style, * p<0.05, ** p<0.01, ***p<0.001, ****p<0.0001. Statistical differences were evaluated using two-way ANOVA with Sidak correction or otherwise stated in figure legends. Graphs were produced and statistical analyses were performed using GraphPad Prism (San Diego, CA, USA). For figure 3 data were compared with the fit of a range of two Bayesian regression models, baseline and interaction.

**Figure 1:**
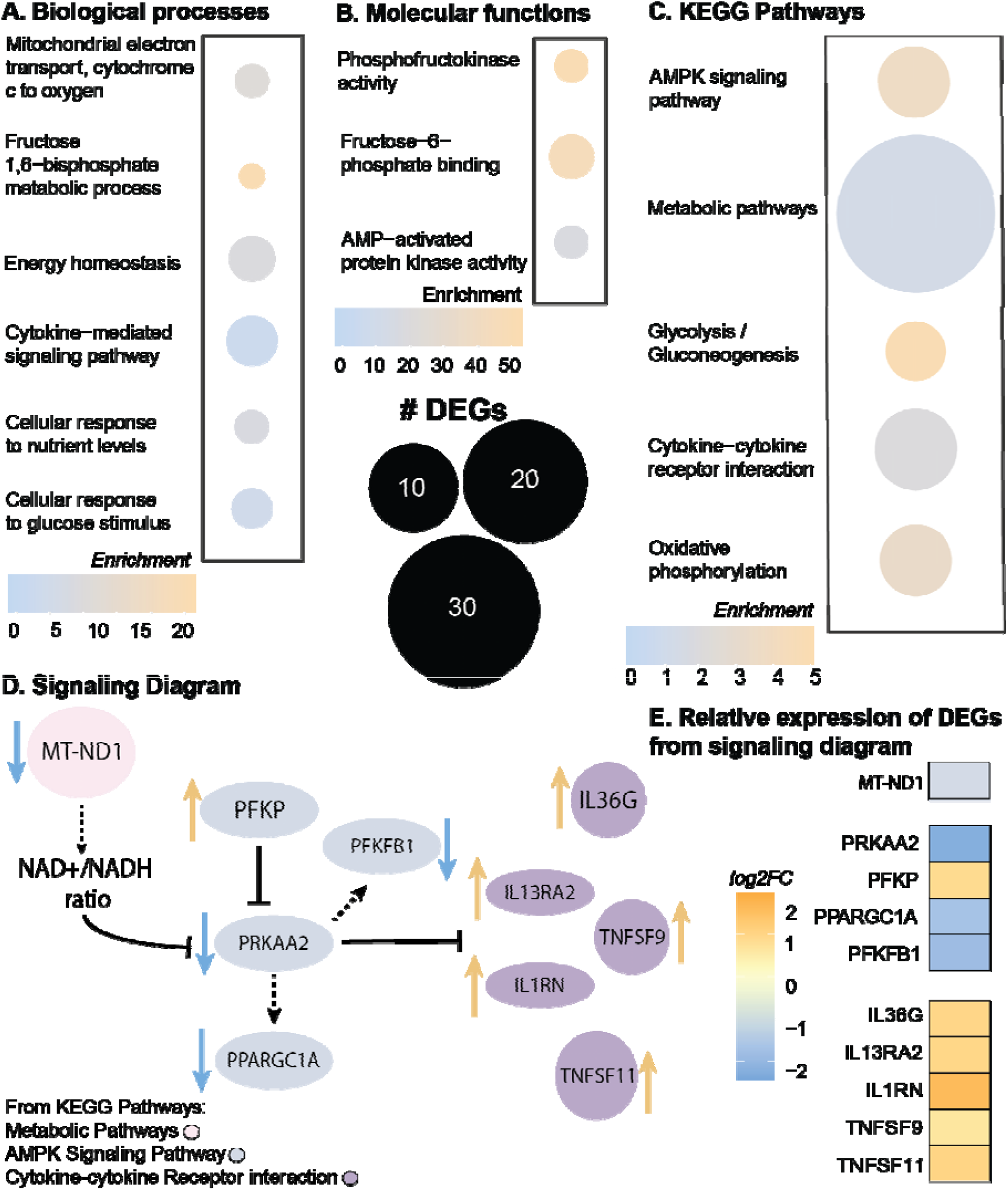
Metabolic pathways and AMPK signaling are dysregulated in human tendinopathy. (A-C) Enrichment quantification of differentially expressed genes (DEGs) from bulk RNA sequencing of human tendinopathic samples compared to healthy control tendons. (A. Enriched GO biological processes; B. Enriched GO molecular functions; C. Enriched Kegg pathways.) (D) Signaling diagram from individual DEGs from KEGG pathways (Gray: DEGs in AMPK signaling pathway; Pink: DEGs from metabolic pathways; Purple: DEGs in cytokine-cytokine receptor interaction pathway; blue arrow representative of downregulated DEGs, orange arrow representative of upregulated DEGs) (E) Relative expression of DEGs from signaling diagram (log2 fold change) from human tendinopathic samples compared to healthy control tendons. n= 7 samples/group. Adjusted p-value: p<0.05, log2FC±1.5

**Figure 2:**
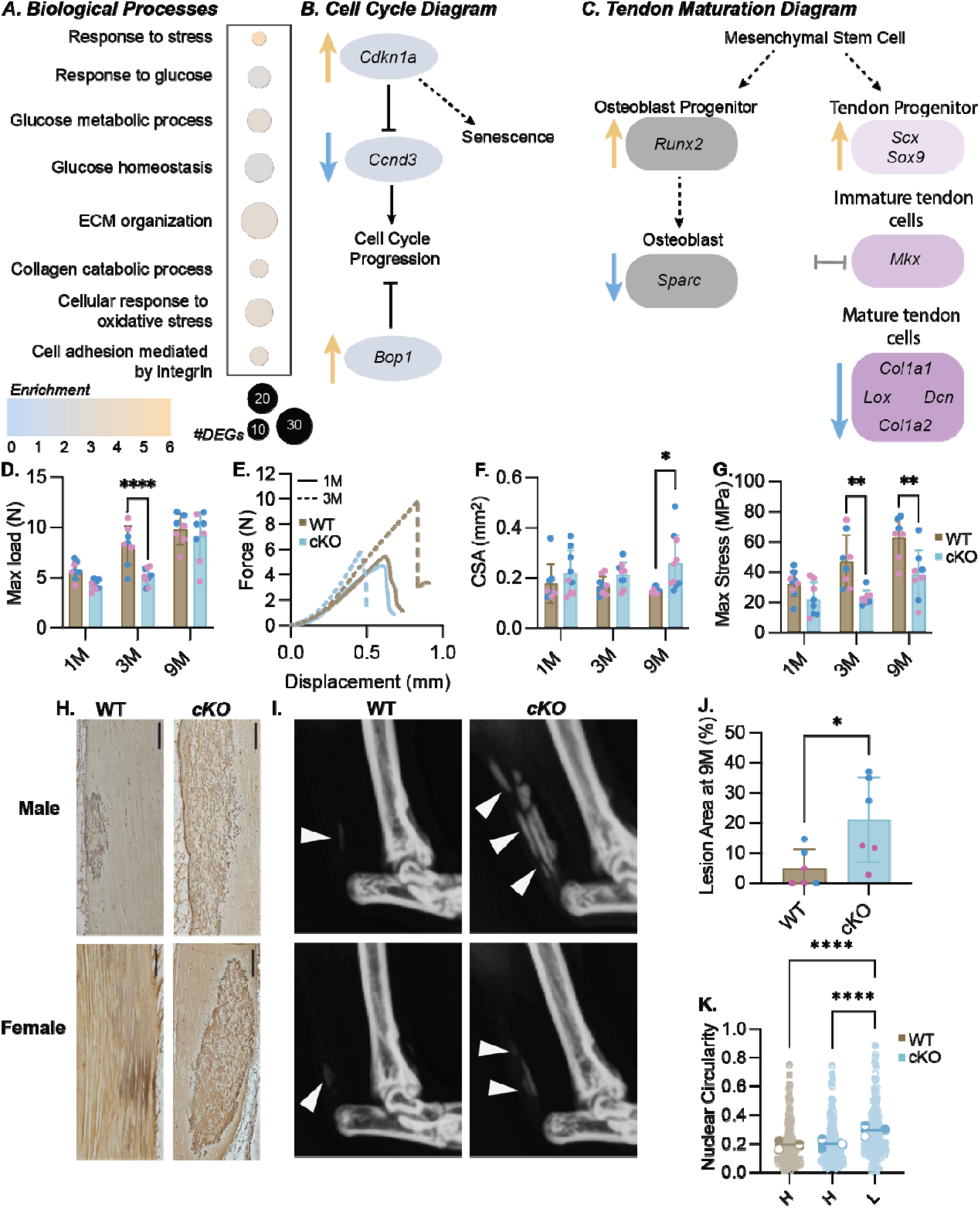
Phenotypic characterization of Prkaa1^Scx-Cre^ cKO Achilles tendons. (A) Quantification of GO biological processes found during bulk RNA sequencing in 1M female Prkaa1^Scx-Cre^ cKO mice compared to WT controls. (B) Signaling diagram created from cell cycle related DEGs in Prkaa1^Scx-Cre^ cKO mice compared WT controls. (blue arrow representative of downregulated DEGs, orange arrow representative of upregulated DEGs) (C) Signaling diagram created from tendon maturation related DEGs in Prkaa1^Scx-Cre^ cKO mice compared WT controls. (blue arrow representative of downregulated DEGs, orange arrow representative of upregulated DEGs, gray bay representative of no significant change) n=3 samples/group. Adjusted p-value: Adjusted p-value: p<.05, log2FC±1.5 (D-G) Biomechanical outcomes from 1, 3 and 9M WT and Prkaa1^Scx-Cre^ cKO Achilles tendons (D. Maximum Load before failure; E. Representative Force vs Displacement curve for 1M and 3M WT and Prkaa1^Scx-Cre^ cKO Achilles tendons F. Cross-sectional area (CSA); G. Maximum stress). n =6 - 10/group, * p<.05, ** p<0.01; two-way ANOVA with Sidak correction. (H) Representative silver nitrate-stained images in 9M WT and Prkaa1^Scx-Cre^ cKO Achilles tendons. Scale bar = 200µm. (I) Representative X-ray images of dystrophic calcification, highlighted by white arrows in the 9M WT and cKO Achilles tendons. (J) Quantification of positive silver nitrate-stained lesions in 9M WT and Prkaa1^Scx-Cre^ cKO Achilles tendons. n =6/group, * p<0.05. (K) Quantification of nuclear circularity 9M WT and Prkaa1^Scx-Cre^ cKO Achilles tendons. ((H) = healthy, (L) = lesion; n =3/group, ****p<.0001; by paired t-test.

**Figure 3:**
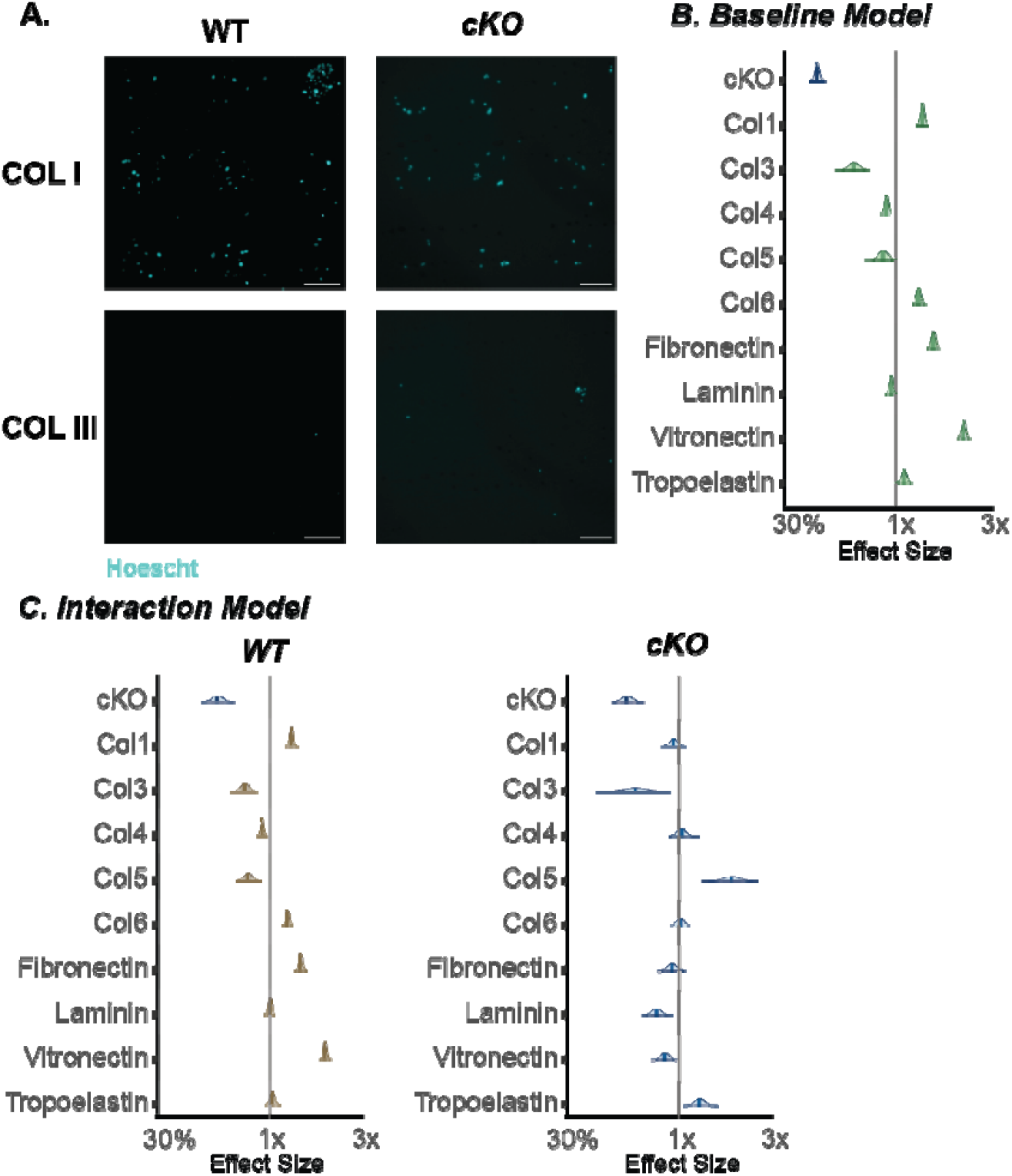
Tendon fibroblasts from Prkaa1^Scx-Cre^ cKO mice have impaired ECM specific adhesion. (A) Representative images of Hoechst staining used for COLI and COL III. (B-C) Posterior distributions with median and 80% confidence intervals of Bayesian regression models quantified from nuclear counts (B. Baseline; C. Interaction; Grey bar represents median number of all nuclear counts regardless of genotype, right side of median indicates high binding affinity, left side of median indicates low binding affinity, green = combined counts, brown=WT, blue= cKO) n = 3/group.

**Figure 4:**
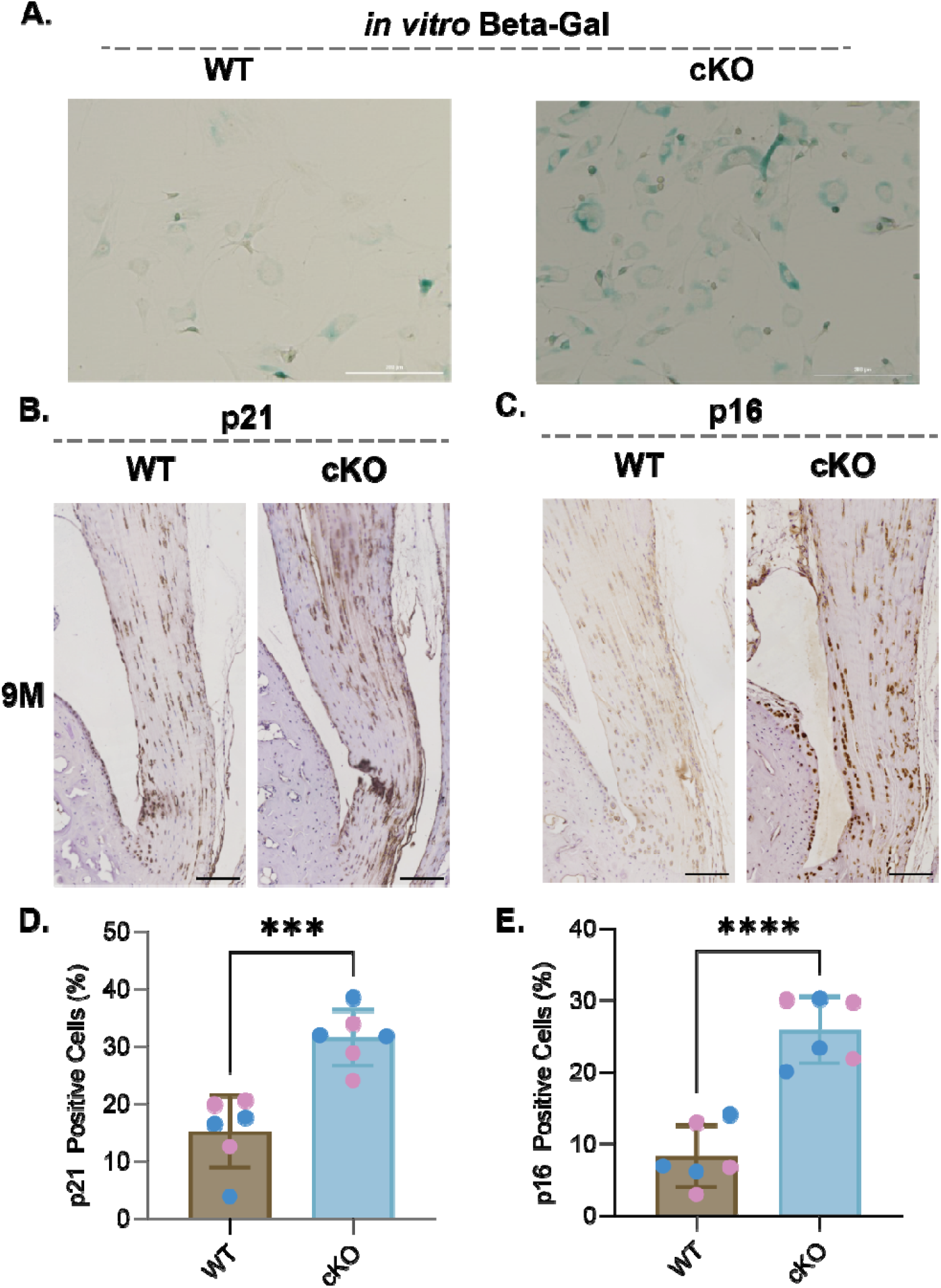
Prkaa1^Scx-Cre^ cKO tendon fibroblasts had increased expression of senescence markers in vitro and in vivo. (A) Representative Beta-Galactosidase stain of WT and Prkaa1^Scx-Cre^ cKO tail TFs *in vitro* after five days in culture. Scale bar = 200µm. n=3/group. (B and C) Representative IHC images in 9M WT and Prkaa1^Scx-Cre^ cKO Achilles tendons of (B) p21 and (C) p16 staining using DAB. (D and E) Quantification of (D) positive p21 cells and (E) positive p16 cells in 9M WT and Prkaa1^Scx-Cre^ cKO Achilles tendons. Scale bar = 500µm. n =6/group (pink = female; blue = male), ***p<0.001, ****p<0.0001; two-way ANOVA with Sidak correction.

**Figure 5:**
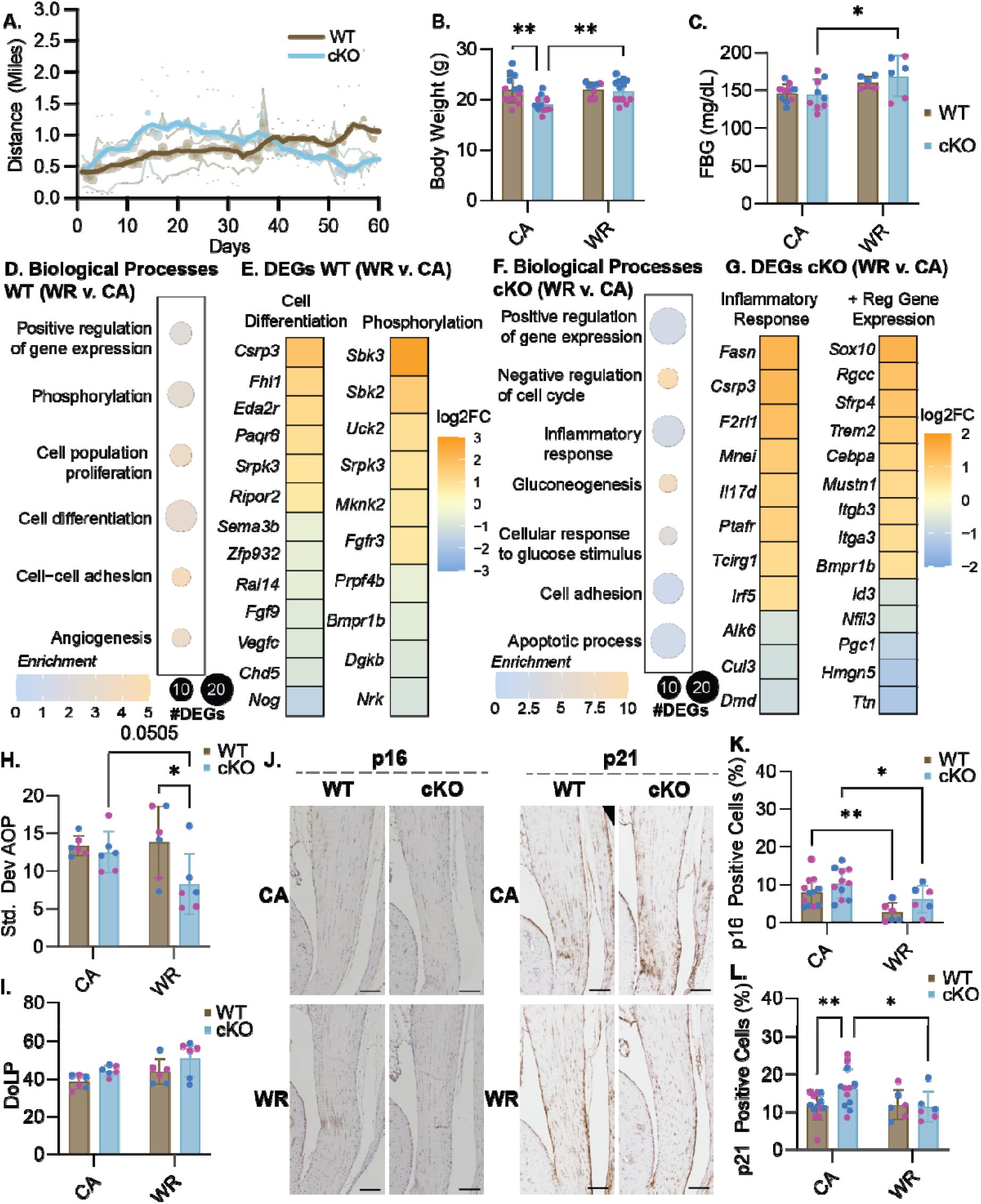
Effects of exercise on Prkaa1^Scx-Cre^ cKO mice. (A) Quantification of distance run for WT and Prkaa1^Scx-Cre^ cKO mice. (B) Quantification of body weight for WT and Prkaa1^Scx-Cre^ cKO WR and CA mice. (C) Quantification of fasting blood glucose (FBG) mg/dL for wheel running groups for WT and Prkaa1^Scx-Cre^ cKO WR and CA mice. n =6-12/group, females = pink; males = blue. * p<0.05, ** p<0.01; two-way ANOVA with Sidak correction. (D) Quantification of GO biological processes found during bulk RNA sequencing in WT WR mice compared to CA controls. (E) Quantification of the log2 fold change of DEGs from WT mice from biological processes WR mice compared to CA controls (Cell differentiation; Phosphorylation). (F) Quantification of GO biological processes found during bulk RNA sequencing in Prkaa1^Scx-Cre^ cKO mice WR mice compared to CA controls. (G) Quantification of the log2 fold change of DEGs in WR Prkaa1^Scx-Cre^ cKO mice from biological processes (Inflammatory response; Positive regulation of gene expression). n= 3-6 samples/group. Adjusted p-value: Adjusted p-value: p<.05, log2FC±1.5 (H and I) Quantification of alignment for WT and Prkaa1^Scx-Cre^ cKO WR and CA mice (H. Degree of linear polarization (DoLP); I. Standard Deviation Angle of Polarization (AOP)), females = pink; males = blue (J) Representative IHC images of p16 and p21 in CA and WR Prkaa1^Scx-Cre^ cKO Achilles tendons. Scale bar = 500µm.Quantification of p16 positive cells. Quantification of positive p21 cells. n =6/group, * p<0.05, ** p<0.001, ***p<0.0001; two-way ANOVA with Sidak correction. (K) Quantification of positive p16 cells. n =6/group, females = pink; males = blue. * p<0.05, ** p<0.01, ***p<0.001; two-way ANOVA with Sidak correction. (L) Quantification of positive p21 cells. n =6/group, females = pink; males = blue. * p<0.05, ** p<0.01, ***p<0.001; two-way ANOVA with Sidak correction.

## RESULTS

### Metabolic pathways and AMPK signaling are dysregulated in tendinopathy

Using bulk RNA-sequencing, we compared healthy hamstring tendon to diseased rotator cuff tendons from human donors and found a downregulation of multiple genes associated with the mitochondrial electron transport chain and energy production, suggesting an overall dysregulation in metabolism (Fig. 1, Table S1 for patient demographics). Enriched biological processes were related to energy homeostasis, response to nutrients, and cytokine signaling (Fig. 1A). Additionally, enriched molecular functions (Fig 1B) and KEGG pathways were also related to AMPK activity and signaling (Fig. 1C), respectively, with KEGG pathways also enriched for regulation of energy metabolism (i.e. glycolysis and oxidative phosphorylation), metabolic pathways and cytokine signaling (Fig. 1C). When analyzing the differentially expressed genes (DEGs) produced from the KEGG pathway results, the AMPK signaling pathway was primarily driven by six DEGs, of which 5 were downregulated in tendinopathic tendons (Fig. 1D, E), with phosphofructokinase (*PFKP*) as the only upregulated gene (Fig. 1D, E). The top 10 hits for DEGs associated with metabolic pathways were also downregulated with tendinopathy, including genes associated with the mitochondrial electron transport chain (Fig. 1D, E). Specifically, tendinopathy samples had downregulated mitochondrial encoded NADH dehydrogenase 1 (*MT-ND1*; Fig. 1D,E), which regulates mitochondrial complex one and the NAD^+^/NADH ratio^30^. A decrease in the NAD^+^/NADH ratio has been associated with decreased AMPK activation and increased production of the senescence associated secretory phenotype (SASP)^31^. Additionally, DEGs associated with the cytokine-cytokine receptor interaction pathway included activation of multiple interleukins (e.g. *IL1, IL13*, and *IL36*; Fig. 1D,E), another characteristic of the SASP^32^. Considering these findings, we sought to identify the role on AMPK in the mouse Achilles tendon in an age-dependent manner.

### Prkaa1^Scx-Cre^ cKO mouse tendons showed dysregulation of cell cycle processes, increased calcification and impaired biomechanical properties

In the Achilles tendons of mice at 1M of age, we identified 621 upregulated and 1008 downregulated genes in Prkaa1^Scx-Cre^ cKO females compared to age-matched WT females using bulk RNA sequencing, yet transcriptional differences were not observed in male mice at 1M or 3M of age. Prkaa1^Scx-Cre^ cKO tendons from female mice at 1M had enriched biological processes related to cell cycle regulation, matrix organization and response to nutrients compared to WT females (Fig. 2A). When comparing cell cycle DEGs, we found a robust upregulation of cell cycle arrest markers in tendons from female Prkaa1^Scx-Cre^ cKO mice such as *Ccngd3*, the gene that encodes cyclin D3, and *Cdkn1a*, the gene that encodes p21 (Fig. 2B). Finally, when comparing metabolism DEGs we identified a significant upregulation of *Bop1*, a known inhibitor of cell cycle progression, in cKO tendons compared to WT at 1M^33^ (Fig.2B). Cellular senescence is defined as terminal growth arrest in cells even if stimulated by mitogens, however they are metabolically and synthetically active^34^. Although there is no clear pathway to senescence, early markers of senescence include changes in glucose metabolism and increased cyclin kinase activity. These findings suggest that Achilles tendons in Prkaa1^Scx-Cre^ cKO mice exhibit a transcriptional profile that supports the onset of senescence^35,36^. Senescent cells are also known to contribute to ECM remodeling ^37^. When comparing tendon and matrix related DEGs, we found *Scx* was significantly upregulated in female Prkaa1^Scx-Cre^ cKO tendons, while *Lox, Loxl3* and both *Col1a1* and *Col1a2* were downregulated compared to WT female tendons (Fig. 2C). A decrease in *Scx* expression in tendon indicates a shift towards a more mature tendon state^38^. Our results showing an increase in Scx mRNA expression in the cKO cells suggest while the WT cells are transitioning into a more mature tendon the cKO cells transcriptionally resemble a progenitor tendon cell. At the protein level, LOX catalyzes the formation of crosslinks in collagen and is essential for the formation of collagen fibril, decreased *Lox* gene expression has been associated shown to reduce the ability to withstand load in *in vitro* tendon constructs ^39^. We found the Prkaa1^Scx-Cre^ cKO Achilles tendons had impaired mechanical function compared to WT controls. First, we found the maximum failure load of Prkaa1^Scx-Cre^ cKO Achilles tendons trended downward at 1M (p = 0.0509) and was significantly reduced at 3M compared to WT controls (Fig. 2D,E), however the maximum load equilibrated by 9M of age (Fig. 2D). Prkaa1^Scx-Cre^ cKO Achilles tendons were significantly larger (cross-sectional area, CSA) compared to WT controls at 9M (Fig. 2F). When normalizing the load to CSA we found peak mechanical stress of Prkaa1^Scx-Cre^ cKO Achilles tendons was also significantly reduced at 3M and 9M of age compared to WT controls (Fig. 2G). When comparing at 3% stain the tangent modulus of Prkaa1^Scx-Cre^ cKO Achilles tendons were significantly reduced at 3M of age (Table S2). Finally, we saw no changes to stiffness between WT and cKO tendons at any age we investigated (Table S2). The functional changes we observed were likely not due to impaired muscle function, as we saw no differences in muscle force production in Prkaa1^Scx-Cre^ cKO mice at 3M (Tables S3-9). We next analyzed the Prkaa1^Scx-Cre^ cKO Achilles tendons histologically and identified positive silver nitrate-stained areas we will report as lesions (Fig.2H), Quantification of silver nitrate-stained lesions at 9M of age showed that Prkaa1^Scx-Cre^ cKO Achilles tendons had a significantly greater percent of positive lesion area compared to WT controls (Fig. 2J) and X-ray confirmed these regions appeared as radio-opaque ectopic calcification (Fig. 2I). Cells contributing to ectopic calcification can exhibit nuclear swelling due to increased stress caused by aberrant calcium levels^40^. In line with this, we found the cell nuclei in lesions at 9M were rounder compared to non-lesion “healthy” tendon within Prkaa1^Scx-Cre^ cKO Achilles tendons as well as in WT tendons (Fig. 2K). Finally, we found that collagen organization, measured using quantitative polarized light imaging (qPLI) of fiber birefringence on histological sections, was not different between Prkaa1^Scx-Cre^ cKO tendons and WT controls at 1, 3 and 9 months of age (Fig. S1A, B). In summary we identified dysregulation of matrix interaction signaling as well as markers of pre-mature senescence therefore, we sought to identify how loss of *Prkaa1* effects ECM specific adhesion and if Prkaa1^Scx-Cre^ cKO tendons exhibited additional senescence markers with age.

### Prkaa1^Scx-Cre^ cKO Achilles tendon fibroblasts have altered ECM specific adhesion and show an increase in senescence associated markers

To compare cellular adhesion between Prkaa1^Scx-Cre^ cKO and WT TFs we plated cells on an array of 36 matrix spots comprised of individual and mixtures of 9 ECM component proteins and measured nuclear counts after 24h as a proxy for adhesion. Across all conditions, spot counts ranged from 0 to 408 with a median of 95. To analyze the drivers of this variation, we fit Bayesian general linear regression models with genotype and matrix proteins as covariates and using weakly informative priors and compared the posterior distributions over regression weights. As a baseline model, we assumed no statistical interaction between genotype and the matrix proteins and since the outcomes are counts, we used a Poisson link function. We found that Prkaa1^Scx-Cre^ cKO TFs were less adherent than WT TFs (Fig. 3A). Additionally, collagen (COL) 1, COL6, fibronectin and vitronectin were more permissive while COL3 was less permissive for adhesion regardless of genotype (Fig. 3A). Considering an interaction model where genotype and matrix interact suggested there was a preference of Prkaa1^Scx-Cre^ cKO TF adhesion for Col5 and tropoelastin. Additionally, Prkaa1^Scx-Cre^ cKO TFs showed a decreased preference for COL1 and laminin (Fig. 3B). There is increasing evidence showing the composition of the ECM potentiates cellular senescence^41^. We identified senescence in Prkaa1^Scx-Cre^ cKO tail TFs using Beta-Galactosidase staining (Fig. 4A). Two cell-cycle inhibitors that are commonly regarded as markers of senescent cells are the cyclin-dependent kinase inhibitors p21 and p16^42^. Here, we used immunohistochemistry of the Achilles tendon and identified a significant increase in p21 and p16 in Prkaa1^Scx-Cre^ cKO mice compared to WT controls at 9M of age (Fig. 4B-E). It is important to note that even in the 9M WT samples we saw >10% of the tendon cell population expressed both p16/p21. Cells expressing senescent markers can play critical roles in tissue regeneration and homeostasis^43^, suggesting that while we saw a significant increase in senescent markers in Prkaa1^Scx-Cre^ cKO Achilles tendons, senescence may be a normal process in tendon homeostasis. Next, we gave WT and Prkaa1^Scx-Cre^ cKO access to voluntary running wheels to explore mechanical and metabolic stress.

### Exercise increases organization and reduces the onset of senescence in Prkaa1^Scx-Cre^ cKO Achilles tendons

We identified a significant linear relationship between distance ran and days with wheels in the WT mice (r^2^=0.7295) during voluntary wheel running (WR) but not the Prkaa1^Scx-Cre^ cKO mice (r^2^=0.02202) showing the WT mice ability to adapt (Fig. 5A). The body weight of Prkaa1^Scx-Cre^ cKO mice was significantly lower than WT mice in cage activity (CA) group but was rescued with WR (Fig. 5B). Additionally, fasting blood glucose (FBG) was significantly elevated in the Prkaa1^Scx-Cre^ cKO WR mice compared to CA mice at 3M of age (Fig. 5C). Voluntary wheel running can influence the metabolic activity in rats where *Gapdh* was shown to be significantly downregulated as well as an increase in overall food intake compared to controls^44^. Using bulk RNA sequencing of Achilles tendons, we found 109 upregulated and 78 downregulated genes with WR compared to CA in WT tendons. In WT tendons, enriched biological processes were related to protein phosphorylation, cell differentiation, and positive regulation of gene expression and angiogenesis (Fig. 5D). When comparing DEGs from cell differentiation processes, we found upregulation of the mechanosensor *Csrp3* (Cysteine and Glycine-rich protein 3) (Fig. 5E). Additionally, when comparing DEGs related to phosphorylation, we saw upregulation of phosphotransferases *Sbk3* and *Sbk2* (SH3 Domain Binding Kinase Family) and downregulation of *Bmpr1b* (Bone Morphogenetic Protein Receptor 1B). Protein phosphorylation, the most common post-translational modification, utilizes ATP and is representative of good energy balance within a cell^45^ suggesting voluntary wheel running promotes energy homeostasis in tendons. In Prkaa1^Scx-Cre^ cKO tendons, we found 169 upregulated and 272 downregulated genes with WR compared to CA. Enriched biological processes included positive regulation of gene expression, inflammatory response, and apoptosis (Fig. 5F). When comparing DEGs from inflammatory response we again found upregulation of *Csrp3* as well as *Cul3* (Cullin 3) a protein involved in the ubiquitin-proteasome system (Fig.5G). When looking at the positive regulation of gene expression we found upregulation of *Itgb3, Itga3* and *Bmpr1b* (Fig.5G). Integrin expression is upregulated with increased mechanical signaling, thought to be necessary to transmit mechanical stimuli to the cells during loading^46^. When looking at the structure of the tendon we found a significant increase in tendon organization in the WR Prkaa1^Scx-Cre^ cKO mice when comparing to the WT WR and a trend upward when comparing to the Prkaa1^Scx-Cre^ cKO CA group (p = 0.0505; Fig. 5H) However, we found no significant change in tendon alignment between the groups (Fig. 5I). Immunohistochemistry of Achilles tendons revealed a significant decrease in senescence-associated p21 with WR compared to CA in Prkaa1^Scx-Cre^ cKO mice (Fig. 5J). Finally, we found WR significantly suppressed senescence-associated p16 regardless of genotype (Fig. 5J).

## DISCUSSION

Defining the role of energy metabolism in the context of tendon health and disease is critical for the advancement of non-surgical therapeutics for patients with tendinopathy. While the role AMPK has been established as a regulator of ECM organization and homeostasis in other musculoskeletal tissues^7–9,19^, the role of AMPK in tendon homeostasis and tendinopathy has previously been unclear. Here we have demonstrated that AMPK signaling is dysregulated in human tendinopathy patients and *Prkaa1* is necessary to maintain homeostasis in adult mouse Achilles tendon. These data demonstrated that loss of *Prkaa1* resulted in premature senescence as well as accelerated onset of ectopic calcification. Additionally, we found that exercise delays the onset of senescence in AMPK deficient cells suggesting exercise may stimulate compensatory pathways that maintain tendon homeostasis.

To the best of our knowledge, we are the first to identify dysregulation of the AMPK signaling pathway in tendinopathy patients. We identified upregulation of genes related to SASP production in tendinopathy patients and these trends are consistent when evaluating other data sets from tendinopathic patients^47^. In a study by Zhu et. al, screening for biomarkers of tendinopathy revealed common processes dysregulated in tendinopathy were related to the cell cycle, mitochondria function and pro-inflammatory responses as well as mTOR and NF-κB signaling pathways^47^. Additionally, other musculoskeletal diseases can be driven by senescence and the SASP^48–50^. In osteoarthritis, it is speculated that synovial macrophages, fibroblasts and osteoclasts promote SASP production, leading to an inflammatory response that enhances disease progression^49,50^. Taken together, our findings, along with previous insights into musculoskeletal disease, suggest progression of tendinopathic disease may be driven by the presence of senescent fibroblasts and maintained through an increased proinflammatory response.

We identified the formation of ectopic calcification in Prkaa1^Scx-Cre^ cKO Achilles tendons, especially males. Surprisingly, our bulk RNA sequencing results showed an upregulation of both *Runx2* and *Dlx5* in female, but not male, Prkaa1^Scx-Cre^ cKO tendons. Runx2 is a key transcription factor that regulates osteoblast differentiation^51^. The relationship between AMPK and Runx2 can vary depending on cell type^51–53^. In osteoblasts, AMPK can regulate Runx2 post-transcriptionally and promote osteogenesis^51^. However, in vascular mesenchymal stem cells, AMPK can downregulate Runx2 expression, and loss of AMPKα1 can promote osteoblastic differentiation^53^. Spatially, we noticed the presence of calcification near the peritenon, suggesting the possibility of cell migration into the tendon core before the onset of calcification. Peritenon cells have been identified as being more stem-like than tendon core cells, expressing several stem cell markers^54^. Additionally, peritenon cells are capable of undergoing osteogenic differentiation *in vitro*^54^. Taken together, TFs and peritenon cells may respond similarly to mesenchymal stem cells with loss of AMPK, pushing them toward an osteogenic state. Future directions will explore the role of the peritenon in the formation of ectopic calcification and how migration and trans-differentiation of these cells may be induced by SASP. The formation of calcification within soft tissues has been linked to increase of senescent cells^16,55,56^. Here we identified loss of *Prkaa1* results in both premature senescence and ectopic calcification in the mouse Achilles tendon. Our data supports proposed mechanisms in which an increase p21 responds to a replicative related process (e.g., DNA damage) which precedes the increase in p16, while both together correspond to late senescent cells and the presence of CDK inhibition and cell cycle arrest^57^. Our data also supports previous findings in tendon suggesting that modulation of the AMPK/mTOR signaling cascade via metformin delays the accumulation of senescent cells, cell morphological changes, and calcification^16,58^. Surprisingly, we found that while we had no sex differences in our senescence associated marker expression, we had distinct differences in calcification area between our male and female mice. Future directions will explore estrogen as a protector of calcification in the female tendon as well as identify the potential of the SASP to induce calcification and additional therapeutics to delay onset of senescence.

Integrins regulate the onset of senescence in fibroblasts^59^ AMPK is a known regulator of β-1 integrin^60^. Consequently, loss of AMPK in fibroblasts can increase cell spreading as well as fibronectin fiber formation^60^. Here, we found transcriptional changes of multiple integrins in both the 1M timepoint as well as with exercise in Prkaa1^Scx-Cre^ cKO tendons. Additionally, we found that TFs from Prkaa1^Scx-Cre^ cKO tails have impaired ECM substrate-specific adhesion. Different ECM substrates have unique integrin binding properties, and taken together these data suggest that AMPK activity is responsive to different ECM substrates and integrin accessibility.

The reduction of senescent cells through exercise has been seen in fibrotic disease models^61,62^. Here we found that voluntary wheel running delayed the onset of senescence markers, p16 and p21, as well as increased cell cycle progression markers (i.e. Regulator of cell cycle progression (*Rgcc*)) in Prkaa1^Scx-Cre^ cKO tendons. In addition to regulating cell cycle progression, *Rgcc* may have protective effects against fibrosis, and stimulation of *Rgcc* correlated with enrichment of ECM organization pathways^63^. These trends correlate with our findings that exercise induced a significant increase in *Rgcc* gene expression as well as increased collagen alignment and integrin expression in Prkaa1^Scx-Cre^ cKO tendons.

In summary, these data show that AMPK, which is dysregulated in tendinopathy, is necessary for the maintenance of homeostasis and mechanical function in the mouse Achilles tendon. Additionally, we found exercise has a protective effect against senescence independent of AMPK and promotes energy homeostasis in tendon. These findings highlight the importance of energy balance metabolism in tendon health and overall improves our understanding of the onset and progression of tendinopathy as well as identification of potential treatment options for tendon disease.

### Limitations of the study

Although our data parallels other bulk-RNA sequencing of human Achilles tendinopathy, our cohort of patients was small with only seven per group focusing on the rotator cuff. Additionally, despite conducting bulk-RNA sequencing at 1M and 3M on female and male mice, we only observed large transcriptional differences in our female 1M mice. We speculate either our timepoint was too late to capture similar differences in the male mice or the phenotype was driven by post translational or other modifications. Future work with proteomics could assist in divulging these differences.

## RESOURCE AVAILABILITY

### Lead contact

Further information and requests for resources and reagents should be directed to and will be fulfilled by the lead contact, Adam Abraham.

### Materials availability

This study did not generate new unique reagents.

### Data and code availability

- Bulk-seq data have been deposited at Gene Expression Omnibus database and are publicly available as of the date of publication.
- All raw data will be deposited to Deep Blue Data

## ACKNOWLEDGMENTS

University of Michigan Center for Cellular Plasticity and Organ Design Emerging Scholars program (ACA); National Science Foundation Graduate Research Fellowship DGE2241144 to LH; National Institutes of Health R01AR079367 to MLK; Michigan Integrative Musculoskeletal Health Core Center P30 AR069620 to MLK; University of Michigan Rackham Graduate Student Research Grant to LH; University of Michigan Integrative Musculoskeletal Health Core Pilot and Feasibility Grant to ACA; National Science Foundation CAREER 144448 to MLK; Library preparation and next-generation sequencing was carried out in the Advanced Genomics Core at the University of Michigan. We are grateful for histological support from Emma Snyder-White and Carol Whitinger (through the Michigan Integrative Musculoskeletal Health Center Core). We are grateful for thoughtful discussions about tendon biomechanics with Sarah Calve at the University of Colorado.

## AUTHOR CONTRIBUTIONS

Conceptualization L.A.H, M.L.K, A.C.A; methodology, L.A.H., M.O., S.L., S.V.B., M.L.K, A.C.A. Investigation, L.A.H., N.M., J.C., S.S.S., P.C., C.D., S-H.B., S.V.B., S.G., S.L., T.P., M.O., M.A., N.A.; writing—original draft, L.A.H., N.M., M.L.K., A.C.A.; writing—review & editing, L.A.H., N.M., J.C., S.S.S., P.C., S-H.B., C.D., S.G.,S.L., T.P., M.O., S.V.B., M.A., N.M., M.L.K., A.C.A; funding acquisition, L.A.H, M.L.K., and A.C.A..; resources, M.L.K. and A.C.A; supervision, M.L.K. and A.C.A.

## SUPPLEMENTAL INFORMATION

**Document S1. Figures S1 and Table S1 -S2**

**Figure S1**. Quantification of alignment for WT and Prkaa1^Scx-Cre^ cKO WR and CA animals, A) degree of linear polarization (DoLP); B) standard deviation of the angle of polarization (AOP).

**Table S1**. Patient Demographics for Bulk-RNA Sequencing

**Table S2**. Biomechanical Properties of Murine Achilles Tendons in wildtype (WT) and *Prkaa1*^*Scx-Cre*^*mice*

**Table S3**. EDL muscle function in 3M wildtype (WT) and *Prkaa1*^*Scx-Cre*^*mice*

**Table S4**. Soleus muscle function in 3M wildtype (WT) and *Prkaa1*^*Scx-Cre*^*mice*

**Table S5**. Gastrocnemius muscle function in 3M wildtype (WT) and *Prkaa1*^*Scx-Cre*^*mice*

**Table S6**. EDL muscle response (frequency sweep) of 3M wildtype (WT) and *Prkaa1*^*Scx-Cre*^*mice*

**Table S7**. Soleus muscle response (frequency sweep) in 3M wildtype (WT) and *Prkaa1*^*Scx-Cre*^*mice*

**Table S8**. Gastrocnemius muscle response to nerve stimulation (frequency sweep) in 3M wildtype (WT) and *Prkaa1*^*Scx-Cre*^*mice*

**Table S9**. Gastrocnemius muscle response with direct muscle stimulation (frequency sweep) in 3M wildtype (WT) and *Prkaa1*^*Scx-Cre*^*mice*

## REFERENCES

1. Winnicki, K., Ochała-Kłos, A., Rutowicz, B., Pękala, P.A., and Tomaszewski, K.A. (2020). Functional anatomy, histology and biomechanics of the human Achilles tendon — A comprehensive review. Annals of Anatomy - Anatomischer Anzeiger 229, 151461. 10.1016/j.aanat.2020.151461.

2. Montagna, C., Svensson, R.B., Bayer, M.L., Rizza, S., Maiani, E., Yeung, C.-Y.C., Filomeni, G., and Kjær, M. (2022). Autophagy guards tendon homeostasis. Cell Death Dis 13, 402. 10.1038/s41419-022-04824-7.

3. Millar, N.L., Silbernagel, K.G., Thorborg, K., Kirwan, P.D., Galatz, L.M., Abrams, G.D., Murrell, G.A.C., McInnes, I.B., and Rodeo, S.A. (2021). Tendinopathy. Nat Rev Dis Primers 7, 1. 10.1038/s41572-020-00234-1.

4. Fouda, M.B., Thankam, F.G., Dilisio, M.F., and Agrawal, D.K. (2017). Alterations in tendon microenvironment in response to mechanical load: potential molecular targets for treatment strategies. Am J Transl Res 9, 4341–4360.

5. Wu, Y.-F., Wang, H.-K., Chang, H.-W., Sun, J., Sun, J.-S., and Chao, Y.-H. (2017). High glucose alters tendon homeostasis through downregulation of the AMPK/Egr1 pathway. Sci Rep 7, 44199. 10.1038/srep44199.

6. Herzig, S., and Shaw, R.J. (2018). AMPK: guardian of metabolism and mitochondrial homeostasis. Nat Rev Mol Cell Biol 19, 121–135. 10.1038/nrm.2017.95.

7. Zong, H., Ren, J.M., Young, L.H., Pypaert, M., Mu, J., Birnbaum, M.J., and Shulman, G.I. (2002). AMP kinase is required for mitochondrial biogenesis in skeletal muscle in response to chronic energy deprivation. Proc. Natl. Acad. Sci. U.S.A. 99, 15983–15987. 10.1073/pnas.252625599.

8. Myers, R.W., Guan, H.-P., Ehrhart, J., Petrov, A., Prahalada, S., Tozzo, E., Yang, X., Kurtz, M.M., Trujillo, M., Gonzalez Trotter, D., et al. (2017). Systemic pan-AMPK activator MK-8722 improves glucose homeostasis but induces cardiac hypertrophy. Science 357, 507–511. 10.1126/science.aah5582.

9. Zhou, S., Lu, W., Chen, L., Ge, Q., Chen, D., Xu, Z., Shi, D., Dai, J., Li, J., Ju, H., et al. (2017). AMPK deficiency in chondrocytes accelerated the progression of instability-induced and ageing-associated osteoarthritis in adult mice. Sci Rep 7, 43245. 10.1038/srep43245.

10. Izumi, S., Otsuru, S., Adachi, N., Akabudike, N., and Enomoto-Iwamoto, M. (2019). Control of glucose metabolism is important in tenogenic differentiation of progenitors derived from human injured tendons. PLoS ONE 14, e0213912. 10.1371/journal.pone.0213912.

11. Patel, S.H., Yue, F., Saw, S.K., Foguth, R., Cannon, J.R., Shannahan, J.H., Kuang, S., Sabbaghi, A., and Carroll, C.C. (2019). Advanced Glycation End-Products Suppress Mitochondrial Function and Proliferative Capacity of Achilles Tendon-Derived Fibroblasts. Sci Rep 9, 12614. 10.1038/s41598-019-49062-8.

12. Ackerman, J.E., Best, K.T., Muscat, S.N., and Loiselle, A.E. (2021). Metabolic Regulation of Tendon Inflammation and Healing Following Injury. Curr Rheumatol Rep 23, 15. 10.1007/s11926-021-00981-4.

13. Grinstein, M., Dingwall, H.L., O’Connor, L.D., Zou, K., Capellini, T.D., and Galloway, J.L. (2019). A distinct transition from cell growth to physiological homeostasis in the tendon. eLife 8, e48689. 10.7554/eLife.48689.

14. White, J.P., Billin, A.N., Campbell, M.E., Russell, A.J., Huffman, K.M., and Kraus, W.E. (2018). The AMPK/p27Kip1 Axis Regulates Autophagy/Apoptosis Decisions in Aged Skeletal Muscle Stem Cells. Stem Cell Reports 11, 425–439. 10.1016/j.stemcr.2018.06.014.

15. McKay, L.K., and White, J.P. (2021). The AMPK/p27Kip1 Pathway as a Novel Target to Promote Autophagy and Resilience in Aged Cells. Cells 10, 1430. 10.3390/cells10061430.

16. Dai, G., Li, Y., Zhang, M., Lu, P., Zhang, Y., Wang, H., Shi, L., Cao, M., Shen, R., and Rui, Y. (2023). The Regulation of the AMPK/mTOR Axis Mitigates Tendon Stem/Progenitor Cell Senescence and Delays Tendon Aging. Stem Cell Rev and Rep 19, 1492–1506. 10.1007/s12015-023-10526-0.

17. Stapleton, D., Mitchelhill, K.I., Gao, G., Widmer, J., Michell, B.J., Teh, T., House, C.M., Fernandez, C.S., Cox, T., Witters, L.A., et al. (1996). Mammalian AMP-activated Protein Kinase Subfamily. Journal of Biological Chemistry 271, 611–614. 10.1074/jbc.271.2.611.

18. Woods, A., Azzout-Marniche, D., Foretz, M., Stein, S.C., Lemarchand, P., Ferré, P., Foufelle, F., and Carling, D. (2000). Characterization of the Role of AMP-Activated Protein Kinase in the Regulation of Glucose-Activated Gene Expression Using Constitutively Active and Dominant Negative Forms of the Kinase. Molecular and Cellular Biology 20, 6704–6711. 10.1128/MCB.20.18.6704-6711.2000.

19. June, R.K., Liu-Bryan, R., Long, F., and Griffin, T.M. (2016). Emerging role of metabolic signaling in synovial joint remodeling and osteoarthritis. Journal Orthopaedic Research 34, 2048–2058. 10.1002/jor.23420.

20. Love, M.I., Huber, W., and Anders, S. (2014). Moderated estimation of fold change and dispersion for RNA-seq data with DESeq2. Genome Biol 15, 550. 10.1186/s13059-014-0550-8.

21. Sherman, B.T., Hao, M., Qiu, J., Jiao, X., Baseler, M.W., Lane, H.C., Imamichi, T., and Chang, W. (2022). DAVID: a web server for functional enrichment analysis and functional annotation of gene lists (2021 update). Nucleic Acids Research 50, W216–W221. 10.1093/nar/gkac194.

22. Huang, D.W., Sherman, B.T., and Lempicki, R.A. (2009). Systematic and integrative analysis of large gene lists using DAVID bioinformatics resources. Nat Protoc 4, 44–57. 10.1038/nprot.2008.211.

23. Martin, M. (2011). Cutadapt removes adapter sequences from high-throughput sequencing reads. EMBnet j. 17, 10. 10.14806/ej.17.1.200.

24. Andrews, S. (2010). FastQC: a quality control tool for high throughput sequence data. https://www.bioinformatics.babraham.ac.uk/projects/fastqc/.

25. Dobin, A., Davis, C.A., Schlesinger, F., Drenkow, J., Zaleski, C., Jha, S., Batut, P., Chaisson, M., and Gingeras, T.R. (2013). STAR: ultrafast universal RNA-seq aligner. Bioinformatics 29, 15–21. 10.1093/bioinformatics/bts635.

26. Li, B., and Dewey, C.N. (2011). RSEM: accurate transcript quantification from RNA-Seq data with or without a reference genome. BMC Bioinformatics 12, 323. 10.1186/1471-2105-12-323.

27. Ewels, P., Magnusson, M., Lundin, S., and Käller, M. (2016). MultiQC: summarize analysis results for multiple tools and samples in a single report. Bioinformatics 32, 3047–3048. 10.1093/bioinformatics/btw354.

28. Bankhead, P., Loughrey, M.B., Fernández, J.A., Dombrowski, Y., McArt, D.G., Dunne, P.D., McQuaid, S., Gray, R.T., Murray, L.J., Coleman, H.G., et al. (2017). QuPath: Open source software for digital pathology image analysis. Sci Rep 7, 16878. 10.1038/s41598-017-17204-5.

29. Weigert, M., and Schmidt, U. (2022). Nuclei instance segmentation and classification in histopathology images with StarDist. 10.48550/ARXIV.2203.02284.

30. Lin, X., Zhou, Y., and Xue, L. (2024). Mitochondrial complex I subunit MT-ND1 mutations affect disease progression. Heliyon 10, e28808. 10.1016/j.heliyon.2024.e28808.

31. Nacarelli, T., Lau, L., Fukumoto, T., Zundell, J., Fatkhutdinov, N., Wu, S., Aird, K.M., Iwasaki, O., Kossenkov, A.V., Schultz, D., et al. (2019). NAD+ metabolism governs the proinflammatory senescence-associated secretome. Nat Cell Biol 21, 397–407. 10.1038/s41556-019-0287-4.

32. Kumari, R., and Jat, P. (2021). Mechanisms of Cellular Senescence: Cell Cycle Arrest and Senescence Associated Secretory Phenotype. Front. Cell Dev. Biol. 9, 645593. 10.3389/fcell.2021.645593.

33. Strezoska, Ž., Pestov, D.G., and Lau, L.F. (2002). Functional Inactivation of the Mouse Nucleolar Protein Bop1 Inhibits Multiple Steps in Pre-rRNA Processing and Blocks Cell Cycle Progression. Journal of Biological Chemistry 277, 29617–29625. 10.1074/jbc.M204381200.

34. Shtutman, M., Chang, B.-D., Schools, G.P., and Broude, E.V. (2017). Cellular Model of p21-Induced Senescence. Methods Mol Biol 1534, 31–39. 10.1007/978-1-4939-6670-7_3.

35. Ogrodnik, M. (2021). Cellular aging beyond cellular senescence: Markers of senescence prior to cell cycle arrest in vitro and in vivo. Aging Cell 20, e13338. 10.1111/acel.13338.

36. Gorgoulis, V., Adams, P.D., Alimonti, A., Bennett, D.C., Bischof, O., Bishop, C., Campisi, J., Collado, M., Evangelou, K., Ferbeyre, G., et al. (2019). Cellular Senescence: Defining a Path Forward. Cell 179, 813–827. 10.1016/j.cell.2019.10.005.

37. Mavrogonatou, E., Pratsinis, H., Papadopoulou, A., Karamanos, N.K., and Kletsas, D. (2019). Extracellular matrix alterations in senescent cells and their significance in tissue homeostasis. Matrix Biology 75–76, 27–42. 10.1016/j.matbio.2017.10.004.

38. Yoshimoto, Y., Takimoto, A., Watanabe, H., Hiraki, Y., Kondoh, G., and Shukunami, C. (2017). Scleraxis is required for maturation of tissue domains for proper integration of the musculoskeletal system. Sci Rep 7, 45010. 10.1038/srep45010.

39. Herchenhan, A., Uhlenbrock, F., Eliasson, P., Weis, M., Eyre, D., Kadler, K.E., Magnusson, S.P., and Kjaer, M. (2015). Lysyl Oxidase Activity Is Required for Ordered Collagen Fibrillogenesis by Tendon Cells. Journal of Biological Chemistry 290, 16440–16450. 10.1074/jbc.M115.641670.

40. Giachelli, C.M. (1999). Ectopic Calcification. The American Journal of Pathology 154, 671–675. 10.1016/S0002-9440(10)65313-8.

41. Blokland, K.E.C., Pouwels, S.D., Schuliga, M., Knight, D.A., and Burgess, J.K. (2020). Regulation of cellular senescence by extracellular matrix during chronic fibrotic diseases. Clinical Science 134, 2681–2706. 10.1042/CS20190893.

42. Campisi, J., and d’Adda Di Fagagna, F. (2007). Cellular senescence: when bad things happen to good cells. Nat Rev Mol Cell Biol 8, 729–740. 10.1038/nrm2233.

43. De Magalhães, J.P. (2024). Cellular senescence in normal physiology. Science 384, 1300–1301. 10.1126/science.adj7050.

44. Legerlotz, K., Schjerling, P., Langberg, H., Brüggemann, G.-P., and Niehoff, A. (2007). The effect of running, strength, and vibration strength training on the mechanical, morphological, and biochemical properties of the Achilles tendon in rats. Journal of Applied Physiology 102, 564–572. 10.1152/japplphysiol.00767.2006.

45. Ardito, F., Giuliani, M., Perrone, D., Troiano, G., and Muzio, L.L. (2017). The crucial role of protein phosphorylation in cell signaling and its use as targeted therapy (Review). International Journal of Molecular Medicine 40, 271–280. 10.3892/ijmm.2017.3036.

46. Popov, C., Burggraf, M., Kreja, L., Ignatius, A., Schieker, M., and Docheva, D. (2015). Mechanical stimulation of human tendon stem/progenitor cells results in upregulation of matrix proteins, integrins and MMPs, and activation of p38 and ERK1/2 kinases. BMC Molecular Biol 16, 6. 10.1186/s12867-015-0036-6.

47. Zhu, Y.X., Huang, J.Q., Ming, Y.Y., Zhuang, Z., and Xia, H. (2021). Screening of key biomarkers of tendinopathy based on bioinformatics and machine learning algorithms. PLoS ONE 16, e0259475. 10.1371/journal.pone.0259475.

48. Falvino, A., Gasperini, B., Cariati, I., Bonanni, R., Chiavoghilefu, A., Gasbarra, E., Botta, A., Tancredi, V., and Tarantino, U. (2024). Cellular Senescence: The Driving Force of Musculoskeletal Diseases. Biomedicines 12, 1948. 10.3390/biomedicines12091948.

49. Zeng, N., Yan, Z.-P., Chen, X.-Y., and Ni, G.-X. (2020). Infrapatellar Fat Pad and Knee Osteoarthritis. Aging and disease 11, 1317. 10.14336/AD.2019.1116.

50. Jeon, O.H., David, N., Campisi, J., and Elisseeff, J.H. (2018). Senescent cells and osteoarthritis: a painful connection. Journal of Clinical Investigation 128, 1229–1237. 10.1172/JCI95147.

51. Wei, J., Shimazu, J., Makinistoglu, M.P., Maurizi, A., Kajimura, D., Zong, H., Takarada, T., Iezaki, T., Pessin, J.E., Hinoi, E., et al. (2015). Glucose Uptake and Runx2 Synergize to Orchestrate Osteoblast Differentiation and Bone Formation. Cell 161, 1576–1591. 10.1016/j.cell.2015.05.029.

52. Chava, S., Chennakesavulu, S., Gayatri, B.M., and Reddy, A.B.M. (2018). A novel phosphorylation by AMP-activated kinase regulates RUNX2 from ubiquitination in osteogenesis over adipogenesis. Cell Death Dis 9, 754. 10.1038/s41419-018-0791-7.

53. Lu, Y., Yuan, T., Min, X., Yuan, Z., and Cai, Z. (2021). AMPK: Potential Therapeutic Target for Vascular Calcification. Front. Cardiovasc. Med. 8, 670222. 10.3389/fcvm.2021.670222.

54. Cadby, J.A., Buehler, E., Godbout, C., Van Weeren, P.R., and Snedeker, J.G. (2014). Differences between the Cell Populations from the Peritenon and the Tendon Core with Regard to Their Potential Implication in Tendon Repair. PLoS ONE 9, e92474. 10.1371/journal.pone.0092474.

55. Dai, G.-C., Wang, H., Ming, Z., Lu, P.-P., Li, Y.-J., Gao, Y.-C., Shi, L., Cheng, Z., Liu, X.-Y., and Rui, Y.-F. (2024). Heterotopic mineralization (ossification or calcification) in aged musculoskeletal soft tissues: A new candidate marker for aging. Ageing Research Reviews 95, 102215. 10.1016/j.arr.2024.102215.

56. Arefin, S., Buchanan, S., Hobson, S., Steinmetz, J., Alsalhi, S., Shiels, P.G., Kublickiene, K., and Stenvinkel, P. (2020). Nrf2 in early vascular ageing: Calcification, senescence and therapy. Clinica Chimica Acta 505, 108–118. 10.1016/j.cca.2020.02.026.

57. Stein, G.H., Drullinger, L.F., Soulard, A., and Dulić, V. (1999). Differential roles for cyclin-dependent kinase inhibitors p21 and p16 in the mechanisms of senescence and differentiation in human fibroblasts. Mol Cell Biol 19, 2109–2117. 10.1128/MCB.19.3.2109.

58. Zhang, J., Li, F., Nie, D., Onishi, K., Hogan, M.V., and Wang, J.H.-C. (2020). Effect of Metformin on Development of Tendinopathy Due to Mechanical Overloading in an Animal Model. Foot Ankle Int. 41, 1455–1465. 10.1177/1071100720966318.

59. Rapisarda, V., Borghesan, M., Miguela, V., Encheva, V., Snijders, A.P., Lujambio, A., and O’Loghlen, A. (2017). Integrin Beta 3 Regulates Cellular Senescence by Activating the TGF-β Pathway. Cell Reports 18, 2480–2493. 10.1016/j.celrep.2017.02.012.

60. Georgiadou, M., Lilja, J., Jacquemet, G., Guzmán, C., Rafaeva, M., Alibert, C., Yan, Y., Sahgal, P., Lerche, M., Manneville, J.-B., et al. (2017). AMPK negatively regulates tensin-dependent integrin activity. Journal of Cell Biology 216, 1107–1121. 10.1083/jcb.201609066.

61. Schafer, M.J., White, T.A., Iijima, K., Haak, A.J., Ligresti, G., Atkinson, E.J., Oberg, A.L., Birch, J., Salmonowicz, H., Zhu, Y., et al. (2017). Cellular senescence mediates fibrotic pulmonary disease. Nat Commun 8, 14532. 10.1038/ncomms14532.

62. Werner, C., Hanhoun, M., Widmann, T., Kazakov, A., Semenov, A., Pöss, J., Bauersachs, J., Thum, T., Pfreundschuh, M., Müller, P., et al. (2008). Effects of Physical Exercise on Myocardial Telomere-Regulating Proteins, Survival Pathways, and Apoptosis. Journal of the American College of Cardiology 52, 470–482. 10.1016/j.jacc.2008.04.034.

63. Luzina, I.G., Rus, V., Lockatell, V., Courneya, J.-P., Hampton, B.S., Fishelevich, R., Misharin, A.V., Todd, N.W., Badea, T.C., Rus, H., et al. (2022). Regulator of Cell Cycle Protein (RGCC/RGC-32) Protects against Pulmonary Fibrosis. Am J Respir Cell Mol Biol 66, 146–157. 10.1165/rcmb.2021-0022OC.

