## Supplemental Data for "AMP-activated protein kinase (AMPK) is essential for tendon homeostasis and prevents premature senescence and ectopic calcification"

**Document S1**

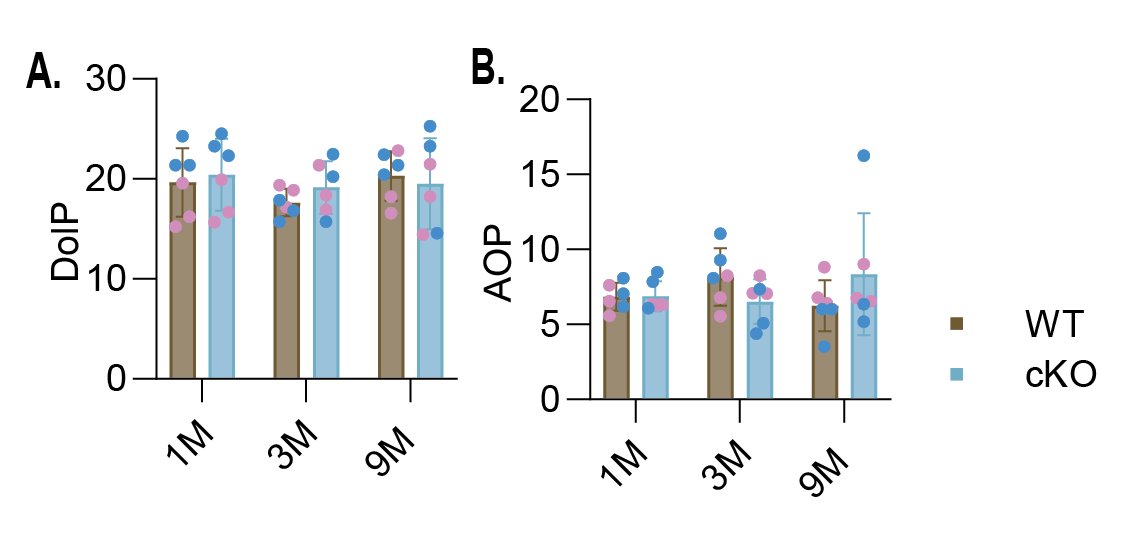

**Figure S1.** Quantification of­­­ alignment for WT and Prkaa1^Scx-Cre^ cKO WR and CA animals, A) degree of linear polarization (DoLP); B) standard deviation of the angle of polarization (AOP).

**Table S1. Patient Demographics for Bulk-RNA Sequencing**

| **Sample ID** | **Condition** | **Age** | **Sex** | **Location** |
| --- | --- | --- | --- | --- |
| 10S | Tendinopathy | 41 | M | Shoulder |
| 13S | Tendinopathy | 54 | M | Shoulder |
| 14S | Tendinopathy | 61 | F | Shoulder |
| 15S | Tendinopathy | 59 | F | Shoulder |
| 16S | Tendinopathy | 59 | M | Shoulder |
| 1T | Normal | 38 | M | Hamstring |
| 2T | Normal | 30 | F | Hamstring |
| 3S | Tendinopathy | 45 | F | Shoulder |
| 3T | Normal | 24 | M | Hamstring |
| 4T | Normal | 29 | M | Hamstring |
| 5T | Normal | 15 | M | Hamstring |
| 6T | Normal | 28 | F | Hamstring |
| 7T | Normal | 22 | F | Hamstring |
| 9S | Tendinopathy | 70 | F | Shoulder |

**Table S2. Biomechanical Properties of Murine Achilles Tendons in wildtype (WT) and *Prkaa1^Scx-Cre^mice***

|  | **WT**  **1M** | **cKO**  **1M** | ***p-*value** | **WT**  **3M** | **cKO**  **3M** | ***p-*value** | **WT**  **9M** | **cKO**  **9M** | ***p-*value** |
| --- | --- | --- | --- | --- | --- | --- | --- | --- | --- |
| Body weight (g) | 16.5 ± 2.18 | 17.73 ± 1.66 | 0.9040 | 23.13 ± 3.07 | 24.52 ± 3.82 | 0.8708 | 27.25 ± 5.64 | 26.28 ± 2.95 | 0.9503 |
| Achilles CSA (mm^2^) | 0.17 ± 0.08 | 0.22 ± 0.09 | 0.5773 | 0.17 ± 0.04 | 0.21 ± 0.05 | 0.5070 | 0.15 ± 0.11 | 0.26 ± .11 | **0.0109** |
| Max. Load (N) | 5.69 ± 0.85 | 4.24 ± 0.53 | 0.0509 | 8.22 ± 1.94 | 5.10 ± 0.84 | **<0.0001** | 9.79 ± 1.554 | 9.14 ± 2.40 | 0.3980 |
| Stiffness (N/mm^2^) | 10.77 ± 1.62 | 8.87 ± 3.43 | 0.6903 | 15.39 ± 3.95 | 13.09 ± 4.35 | 0.5292 | 18.39 ± 3.24 | 22.98 ± 5.10 | 0.0715 |
| Max. Stress (MPa) | 32. 12 ± 7.92 | 21.87 ± 10.45 | 0.2824 | 47.09 ± 16.18 | 22.87 ± 4.51 | **0.0013** | 62.82 ± 12.83 | 37.96 ± 15.5 | **0.0015** |
| Tan. Mod. (MPa) at 3% Strain | 307.68 ±  65.01 | 175.52 ± 54.11 | 0.5475 | 385.1 ± 215.5 | 247.73 ± 169.4 | **0.0438** | 493.80 ± 111.7 | 482.37 ± 141.7 | 0.9986 |

Values are means ± SD; n =6-9 per group. CSA = Cross-sectional area; Tan. Mod. = Tangent Modulus. Comparisons made vs. WT controls at same age using two-way ANOVA with Sidak correction.

**Table S3. EDL muscle function in 3M wildtype (WT) and *Prkaa1^Scx-Cre^mice***

| **Group** | **Sex** | **Muscle mass (mg)** | **Lo (mm)** | **Lf (mm)** | **CSA (mm^2^)** | **Po (mN)** | **sPo (kN/m^2^)** | **Post-stretch Po (mN)** | **Force Deficit** |
| --- | --- | --- | --- | --- | --- | --- | --- | --- | --- |
| WT | M | 9.9 ± 1.4 | 12.7 ± 0.7 | 5.57 ± 0.30 | 1.67 ± 0.16 | 397 ± 25 | 240 ± 32 | 368 ± 17* | 10% ± 2 |
| WT | F | 7.2 ± 0.5 | 11.7 ± 0.6 | 5.13 + 0.27 | 1.32 ± 0.03 | 286 ± 36 | 216 ± 22 | 246 ± 6* | 8% ± 0* |
| cKO | M | 8.5 ± 0.9 | 11.4 ± 0.6 | 5.00 ± 0.28 | 1.60 ± 0.07 | 374 ± 51 | 233 ± 25 | 328 ± 41 | 12% ± 1 |
| cKO | F | 5.7 ± 1.1 | 11.0 ± 0.3 | 4.85 ± 0.13 | 1.10 ± 0.2 | 256 ± 55 | 232 ± 7 | 236 ± 42 | 7% ± 4 |

Values are means ± SD; n = 3 per group, unless labeled with * (* indicates n=2 per group).

**Table S4. Soleus muscle function in 3M wildtype (WT) and *Prkaa1^Scx-Cre^mice***

| **Group** | **Sex** | **Muscle mass (mg)** | **Lo (mm)** | **Lf (mm)** | **CSA (mm^2^)** | **Po (mN)** | **sPo (kN/m^2^)** | **Post-stretch Po (mN)** | **Force Deficit** |
| --- | --- | --- | --- | --- | --- | --- | --- | --- | --- |
| WT | M | 7.4 ± 0.9 | 10.8 ± 0.3 | 7.64 ± 0.23 | 0.92 ± 0.09 | 216 ± 50 | 234 ± 32 | 195 ± 49 | 10.2% ± 2.1 |
| WT | F | 5.5 ± 0.5 | 10.3 ± 0.3 | 7.31 ± 0.25 | 0.71 ± 0.09 | 166 ± 19 | 235 ± 36 | 146 ± 18 | 12.2% ± 2.1 |
| cKO | M | 6.7 ± 1.1 | 11.2 ± 0.9 | 7.95 ± 0.61 | 0.79 ± 0.07 | 204 ± 20 | 258 ± 8 | 185 ± 24 | 9.6% ± 3.0 |
| cKO | F | 4.7 ± 1.4 | 10.4 ± 0.6 | 7.41 ± 0.39 | 0.59 ± 0.15 | 152 ± 35 | 257 ± 8 | 132 ± 31 | 13.1% ± 1.5 |

Values are means ± SD; n =3 per group.

**Table S5. Gastrocnemius muscle function in 3M wildtype (WT) and *Prkaa1^Scx-Cre^mice***

| **Group** | **Sex** | **Muscle mass (mg)** | **Lo (mm)** | **Lf (mm)** | **CSA (mm^2^)** | **Pt (mN)** | **Po freq (Hz)** | **Po-nerve (mN)** | **sPo-nerve (N/cm^2^)** |
| --- | --- | --- | --- | --- | --- | --- | --- | --- | --- |
| WT | M | 138.6 ± 13.8 | 15.1 ± 0.3 | 6.78 ± 0.14 | 19.27 ± 1.77 | 951 ± 101 | 207 ± 23 | 4710 ± 344 | 24.47 ± 0.83 |
| WT | F | 99.9 ± 0.8 | 14.4 ± 0.4 | 6.48 ± 0.16 | 14.55 ± 0.26 | 708 ± 137 | 187 ± 12 | 3296 ± 189 | 22.64 ± 1.02 |
| cKO | M | 143.7 ± 15.1 | 15.2 ± 1.0 | 6.83 ± 0.45 | 19.84 ± 1.00 | 1050 ± 88 | 193 ± 23 | 4401 ± 310 | 22.27 ± 2.69 |
| cKO | F | 90.6 ± 17.0 | 14.6 ± 0.7* | 6.57 ± 0.32* | 13.44 ± 2.59* | 713 ± 396* | 200 ± 28* | 3193 ± 851* | 23.58 ± 1.79* |

Values are means ± SD; n =3 per group, unless labeled with * (* indicates n=2 per group).

**Table S6. EDL muscle response (frequency sweep) of 3M wildtype (WT) and *Prkaa1^Scx-Cre^mice***

| **Group** | **Sex** | **40Hz** | **80Hz** | **120Hz** | **160Hz** | **180Hz** | **200Hz** | **220Hz** |
| --- | --- | --- | --- | --- | --- | --- | --- | --- |
| WT | M | 246 ± 40 | 356 ± 32 | 386 ± 29 | 395 ±27 | 395 ± 26 | 383 ± 7* | 300 ± 6* |
| WT | F | 178 ± 37 | 253 ± 42 | 273 ± 42 | 281 ± 41 | 283 ± 39 | 284 ± 37 | 265 ± 7* |
| cKO | M | 185 ± 39 | 329 ± 44 | 363 ± 46 | 371 ±47 | 369 ± 48 | 363 ± 73* | NA |
| cKO | F | 139 ± 33 | 217 ± 56 | 241 ± 60 | 250 ± 58 | 252 ± 57 | 254 ± 58 | 255 ± 56 |

Values in mN and are means ± SD; n = 3 per group, unless labeled with * (* indicates n=2 per group).

**Table S7. Soleus muscle response (frequency sweep) in 3M wildtype (WT) and *Prkaa1^Scx-Cre^mice***

| **Group** | **Sex** | **40Hz** | **80Hz** | **120Hz** | **160Hz** | **180Hz** | **200Hz** | **220Hz** |
| --- | --- | --- | --- | --- | --- | --- | --- | --- |
| WT | M | 182 ± 37 | 206 ± 45 | 214 ± 48 | 215 ± 49 | 215 ± 49 | 217 ± 71* | NA |
| WT | F | 141 ± 13 | 156 ± 15 | 161 ± 17 | 163 ± 18 | 165 ± 19 | 166 ± 18 | 172 ± 23* |
| cKO | M | 170 ± 8 | 193 ± 13 | 200 ± 16 | 202 ± 18 | 202 ± 19 | 207 ± 25* | 208 ± 27* |
| cKO | F | 132 ± 31 | 145 ± 35 | 149 ± 36 | 151 ± 35 | 151 ± 35 | 151 ± 35 | NA |

Values in mN and are means ± SD; n =3 per group, unless labeled with * (* indicates n=2 per group).

**Table S8. Gastrocnemius muscle response to nerve stimulation (frequency sweep) in 3M wildtype (WT) and *Prkaa1^Scx-Cre^mice***

| **Group** | **Sex** | **40Hz** | **80Hz** | **120Hz** | **160Hz** | **180Hz** | **200Hz** | **220Hz** |
| --- | --- | --- | --- | --- | --- | --- | --- | --- |
| WT | M | 1508 ± 49 | 3531 ± 250 | 4283 ± 252 | 4533 ± 299 | 4641 ± 274 | 4657 ± 304 | 4892 ± 193 |
| WT | F | 1224 ± 349 | 2638 ± 473 | 3069 ± 352 | 3201 ± 282 | 3250 ± 267 | 3289 ± 183 | NA |
| cKO | M | 1561 ± 216 | 3699 ± 56 | 4198 ± 172 | 4336 ± 273 | 4384 ± 332 | 4348 ± 343 | 4413 ± 397* |
| cKO* | F | 1129 ± 629* | 2435 ± 1051* | 2842 ± 1105* | 3002 ± 1046* | 3085 ± 1005* | 3127 ± 843* | NA |

Values in mN and are means ± SD; n =3 per group, unless labeled with * (* indicates n=2 per group).

**Table S9. Gastrocnemius muscle response with direct muscle stimulation (frequency sweep) in 3M wildtype (WT) and *Prkaa1^Scx-Cre^mice***

| **Group** | **Sex** | **40Hz** | **80Hz** | **120Hz** | **160Hz** | **180Hz** | **200Hz** | **220Hz** |
| --- | --- | --- | --- | --- | --- | --- | --- | --- |
| WT | M | 1208 ± 142 | 3758 ± 342 | 4429 ± 481 | 4602 ± 476 | 4636 ± 444 | 4604 ± 456 | NA |
| WT | F | 1342 ± 608 | 2644 ± 148 | 3019 ± 149 | 3110 ± 179 | 3121 ± 138 | 3109 ± 168 | NA |
| cKO | M | 1621± 254 | 3777 ± 318 | 4138 ± 305 | 4317 ± 228 | 4240 ± 203 | NA | NA |
| cKO* | F | 1217 ± 439* | 2554 ± 539* | 2969 ± 518* | 3119 ± 484* | 3145 ± 459* | 3169 ± 441* | 3110 ± 399* |

Values in mN and are means ± SD; n =3 per group, unless labeled with * (* indicates n=2 per group).
